# Annexin A1, Calreticulin and High Mobility Group Box 1 are elevated in Secondary Progressive Multiple Sclerosis: Does Immunogenic Cell Death Occur in Multiple Sclerosis?

**DOI:** 10.1101/2024.01.12.575470

**Authors:** Mohammad Saeid Hejazi, Sevda Jafari, Soheila Montazersaheb, Ommoleila Molavi, Vahid Hoseini, Mahnaz Talebi, Masoud Nikanfar

**Affiliations:** Molecular Medicine Research Center, Tabriz University of Medical Sciences, Tabriz, Iran; Department of Pharmaceutical Biotechnology, Faculty of Pharmacy, Tabriz University of Medical Sciences, Tabriz, Iran; Nutrition Research Center, Tabriz University of Medical Sciences, Tabriz, Iran; Neuroscience Research Center, Tabriz University of Medical Sciences, Tabriz, Iran; Razi Hospital, Tabriz University of Medical Sciences, Tabriz, Iran

**Keywords:** Multiple sclerosis, Immune response, DAMPs, Inflammatory response, ICD

## Abstract

Multiple sclerosis (MS) is a chronic neuroinflammatory diseases characterized by demyelination of the nerve fibers. Immunogenic cell death (ICD) is a process, during which damaged and stressed cells release danger-associated molecular patterns (DAMPs) activating immune responses. This study aimed to elucidate the induction of ICD in MS diseases. To achieve this goal, the level of DAMPs including Annexin A1, calreticulin and HMGB1 was measured in the cerebrospinal fluid (CSF) of a secondary progressive multiple sclerosis (SPMS) patient in comparison to control group. Results showed significant upregulation (more than two-fold) of Annexin A1, calreticulin and HMGB1 in the CSF of the patient. Although further studies are suggested in this regard, this data could imply induction of ICD in MS. The proposed ICD might trigger immune response against neural cells resulting in neuroinflammation and demyelination in CNS in MS. Our observation could suggest inclusion of ICD interfering treatments in routine MS therapy.

## Introduction

Multiple sclerosis (MS) is characterized by demyelination of the central nervous system (CNS) ^1^. Although, various environmental and genetic factors are involved in MS onset, the role of Epstein-Barr Virus (EBV) infection is approved as an environmental causal agent triggering the disease. ^2,3^ Approximately 2.5 million people worldwide are affected by this disease, which causes substantial neurological disabilities in young adults. ^1^ It has been shown that infiltration of immune cells from the periphery into the CNS results in localized inflammation, demyelination, and axonal damage. Clinical manifestations of MS include cognitive deficits, sensory abnormalities, paralysis, and ocular symptoms that correlate with relapse and remission of the disease. However, the symptoms vary depending on the affected area within the CNS. The disorder manifests in four distinct phenotypes based on its progression and course: relapsing-remitting MS (RRMS), secondary progressive MS (SPMS), primary progressive MS (PPMS), and progressive-remitting MS (PRMS). ^4,5^

Researchers are currently focused on two paradigms, the “outside-in” and “inside-out” to explore the basic etiology and pathophysiological theory of MS. Each paradigm is studied by distinct experimental animal models. The “outside-in” model suggests that MS originates from peripherally-induced inflammatory and autoimmune attacks targeting damaged myelin. Conversely, the “inside-out” theory proposes that primary cell degeneration, particularly in oligodendrocytes (OLs) within the CNS, elicits subsequent reactive inflammatory/autoimmune reactions against myelin fragments. ^6-9^ In this regard, a concept involving a cell death model and a positive feedback loop driven by damage-associated molecular patterns (DAMPs) has been proposed for chronic inflammatory demyelination. This paradigm aims to align with the “inside-out” theory of MS.

Immunogenic cell death (ICD) is a process by which dying cells release danger signals that can activate both innate and adaptive immune responses. ICD occurs in response to infection following the release of pathogen-associated molecular patterns (PAMPs) and also happens in a sterile environment next to the release of danger-associated molecular patterns (DAMPs). DAMPs are endogenous molecules considered as danger signals capable of eliciting the innate and adaptive immune responses, thereby enhancing overt autoimmune disorders. Therefore, ICD is considered not only as a defense against infection, but also a mechanism for clearing damaged and stressed cells. ^10,11^ICD is a subset of regulated cell death (RCD) triggering immune responses. ^12,13^Various types of cell death such as apoptosis, pyroptosis, autophagy-dependent cell death, necroptosis and ferroptosis have been identified as significant contributors to the development of neuroinflammatory and neurodegenerative disorders in the CNS. ^14^ Environmental factors such as EBV infections, smoking and UVR involved in 70% of all autoimmune diseases have been shown to promote forms of cell death leading to the induction of DAMPs. ^13,15^ One instance involves microglia activation triggered by DAMPs like extracellular ATP, which have been observed to intensify neuroinflammation in conditions like Alzheimer’s disease. ^16,17^ Additionally, it is shown that heat shock protein 70 (HSP70) is upregulated in MS ^18^ and high mobility group box 1 (HMGB1) can induce experimental autoimmune encephalomyelitis EAE. ^19^ Compelling evidence suggesting oxidative stress results in the release of specific DAMPs, such as calreticulin (CRT), and cell death leading to neurodegeneration in MS. ^20,21^ To date, there have been no direct reports regarding ICD induction in the CNS of individuals diagnosed with MS.

We hypothesized there are various reasons for the incidence of ICD in MS, including EBV infection. EBV infection is approved as a main causal agent of MS. ^3^ Taking into account the elevated levels of EBV miRNAs found in the CSF exosomes of RRMS patients, implying variation in the virus activity in these patients ^22,23^ and considering that infection could trigger ICD, ^10^ in the present case study we aimed to answer this question: Does ICD happen in MS patients? In order to answer the question, we measured the level of DAMPs, including CRT, Annexin A1 and HMGB1 in the cerebrospinal fluid (CSF) of the SPMS patient compared to the control group.

## Materials and Methods

### Participants and Human Ethics

All participants underwent examination at Razi Hospital in 2023. The secondary progressive multiple sclerosis (SPMS) patient met the McDonald’s criteria ^24^ and the diagnosis was fulfilled in agreement with the recent diagnostic criteria. While the clinical differentiation between RRMS and progressive forms remains challenging, the revised “Lublin Criteria were used to distinguish these phenotypes. ^25^ The case study consisted a total of four individuals: one with SPMS, and three with idiopathic intracranial hypertension (IIH), who served as the control group. The relevant clinical details and demographic information of the participants are presented in Table 1. The current study was approved by the ethics committee of Tabriz University of Medical Sciences with IR.TBZMED.REC.1402.348 number. Written informed consent was obtained from the SPMS patient. The IIH samples were collected from the hospital, where the IIH patients were lumber punctured as a routine treatment.

**Table 1.**
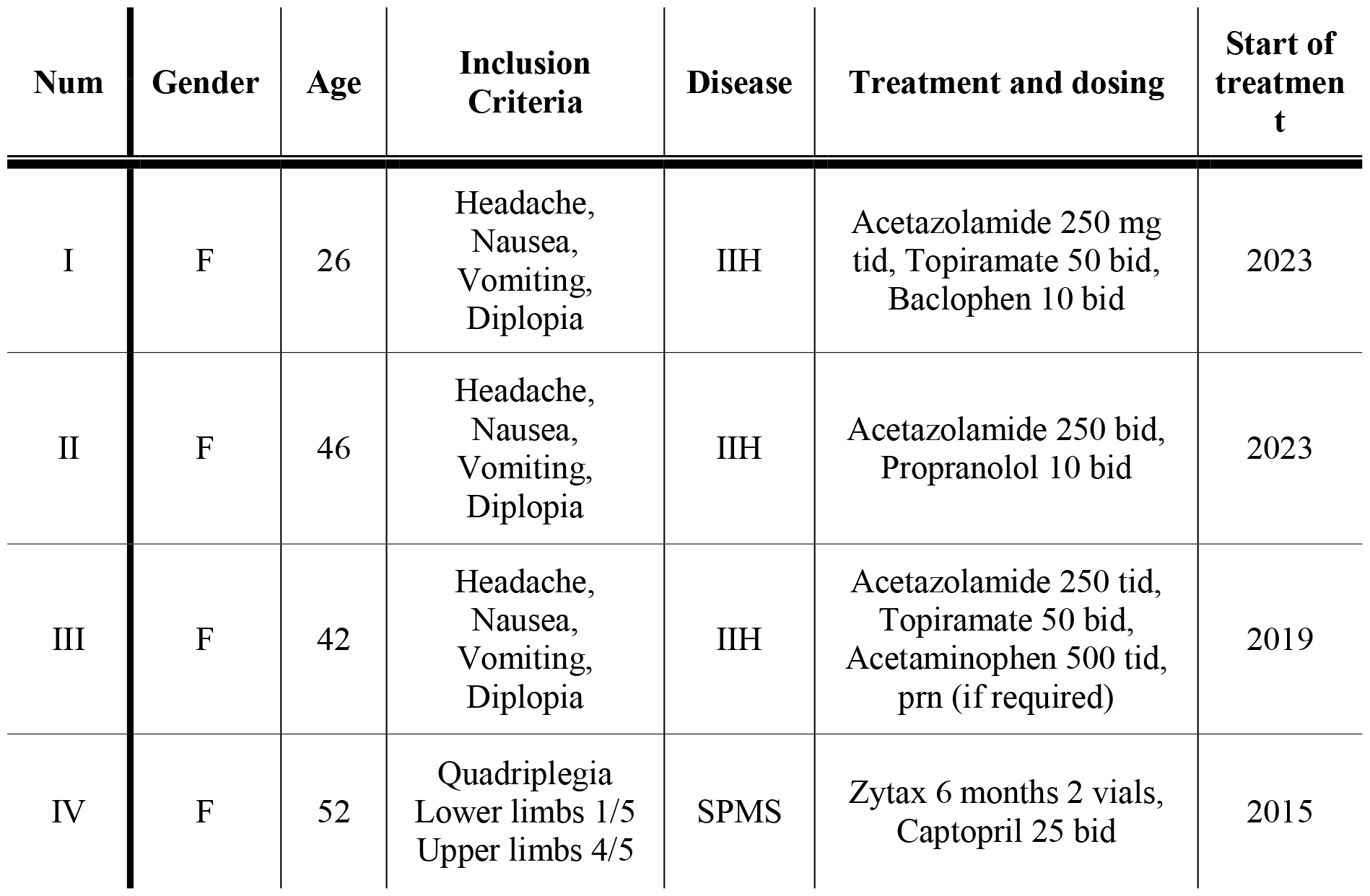
List of the participants enrolled in this study (n=4). F: female; IIH: idiopathic intracranial hypertension; SPMS: secondary progressive MS; tds: three times per day; bid: two times per day, prn: as needed.

### CSF Preparation

CSF samples were collected from IIH individuals and SPMS patient without the need for additional LP. In other words, the remaining CSF samples were collected from the hospital and used in this study. The patient and the healthy control groups were matched in terms of sex and ethnicity.

### Determination of the expression level DAMPs

An Enzyme-linked immunosorbent assay (ELISA) was used to determine the expression levels of DAMP molecules, namely CRT, Annexin A1 and HMGB1. CSF samples were obtained from all participants by non-traumatic lumbar puncture and stored at -80°C until being used. Human ANXA1 commercial ELISA Kit (Elabscience; E-EL-H5512) was used to determine the level of Annexin A1. HMGB1 levels were determined using commercial Novus ELISA kit (cat. No. NBP2-62766). CRT levels were analyzed using Cusabio ELISA kit (Cat. No. CSB- E09787h). The DAMPs levels were standardized based on the protein concentration of each sample. All samples were analyzed in triplicate.

### Statistics

All values are expressed as the mean ± SD. GraphPad Prism Version 9 (GraphPad Software, San Diego, Calif. USA) was used to analyze data. The data from the two groups was compared by t test. The significance level was set at p≤ 0.05.

## Results and Discussion

Our results reveal a significant elevation in the level of Annexin A1 in the CSF of SPMS patient compared to individuals without MS, as depicted in Fig.1. The observed increase in Annexin A1 levels in the CSF of the patient suggests a potential association between Annexin A1 and the pathophysiology of MS. Annexin A1 is known for its involvement in immunogenic cell death processes. It is involved in the functional maturation of antigen-presenting cells (APCs), particularly dendritic cells (DCs) during ICD. Upon release from apoptotic cells, Annexin A1 binds to the formyl peptide receptor 1 receptor (FPR1) on APCs, facilitating a stable interaction between the APC and the dying cancer cell. This function of Annexin A1 enables the uptake of antigens and their cross-presentation on major histocompatibility complex class I (MHC I), ultimately leading to the activation of immune responses. ^26,27^

**Fig. 1.**
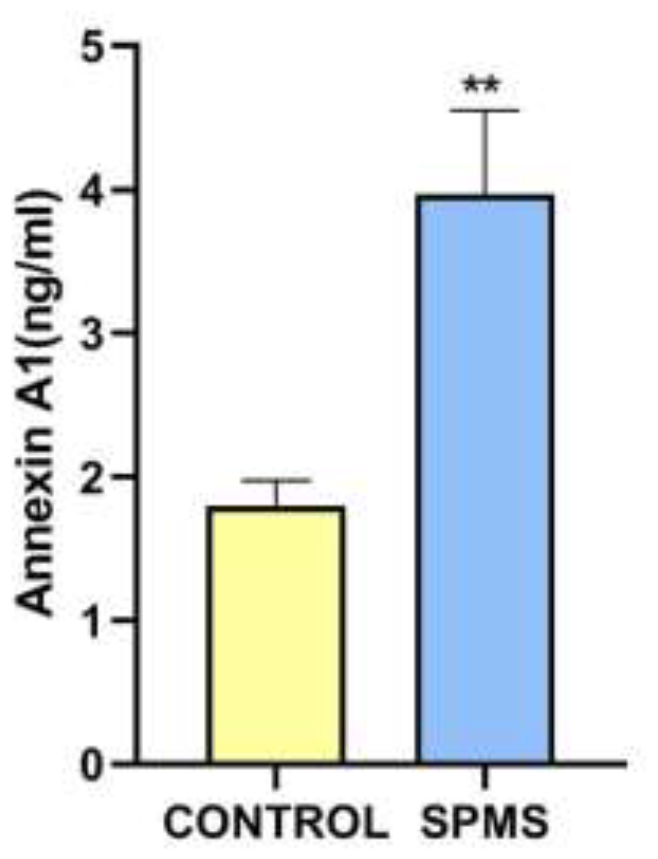
Annexin A1 level in CSF sample of healthy individuals and SPMS patient. Bar graphs related to Annexin A1 measurement using standard ELISA assay that represent means ± SD measured in each group. A comparison between the results from healthy individuals and SPMS patient was done using a t-test. ELISA experiments were repeated at least three times. ** indicates: p < 0.01.

We also identified a significant increase in the level of CRT in the CSF of the patient compared to the levels observed in the control group (Fig.2). CRT, a chaperone protein located in the endoplasmic reticulum (ER), is crucial for proper protein folding, and disruptions in this process can lead to ER stress, implicated in neurodegenerative disorders like Alzheimer’s and Parkinson’s diseases. ^28^ CRT is linked to apoptotic pathways and influences cell death in neurodegenerative conditions. Consistent with our findings, externalized CRT signals phagocytic cells to clear damaged cells, contributing to the inflammation observed in various neurological disorders and some autoimmune diseases. ^20,28-31^ CRT, a multifunctional protein, serves as an “eat-me” signal when exposed on dying cells, aiding in phagocytic clearance and providing insights into ICD-related molecular events. ^26,27^ Consistent with our findings, earlier studies have documented an upregulation of molecules associated with the endoplasmic reticulum stress-related signaling pathway, including CRT, in MS lesions. ^32^ As a multifunctional protein involved in various cellular processes, CRT has also been implicated in ICD and immune responses. The elevated CRT levels may reflect ongoing immunogenic cellular stress or damage associated with neurodegenerative processes in the central nervous system in MS. Further exploration of the specific implications and mechanisms related to increased CRT levels could deepen our understanding of disease progression and potentially inform targeted therapeutic strategies.

**Fig. 2.**
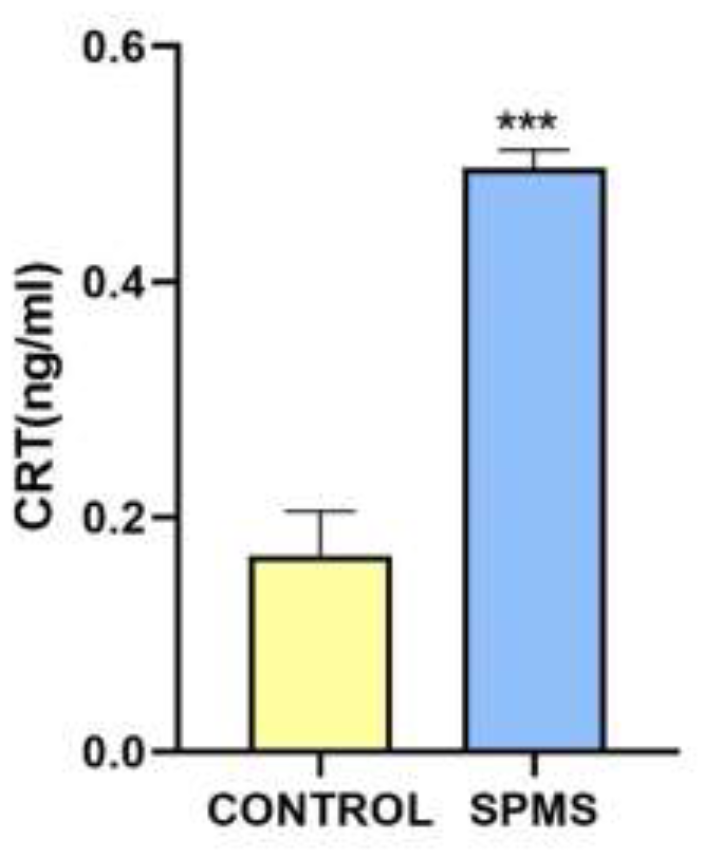
CSF concentration of Calreticulin in healthy individuals and SPMS patient. Calreticulin levels were measured using ELISA assay. The bar graph indicates the level of calreticulin in healthy controls and SPMS patient. Results showed a statistically significant difference between healthy controls and SPMS patient (p>0.001). ELISA experiments were repeated at least three times. *** indicates: p < 0.001.

As shown in Fig. 3. the level of HMGB1 was also significantly elevated in the CSF of the SPMS patient compared to the levels observed in the control group. HMGB1 is a protein crucial in regulating inflammation and immune responses, released during cell injury to signal the immune system. It interacts with receptors like Toll-like receptors and RAGE, triggering inflammatory responses. Previous studies have noted the involvement of HMGB1 in neuroinflammation, contributing to immune responses in neurodegenerative diseases, stroke, traumatic brain injury, epilepsy, and neurodegenerative disorders like Alzheimer’s and Parkinson’s diseases. ^33-35^ Our finding underscores the potential involvement of HMGB1 in the pathogenesis of multiple sclerosis. HMGB1, a nuclear protein with diverse functions, is known for its role in inflammatory responses and cell death processes. ^36,37^ The heightened levels of HMGB1 may reflect ongoing cellular stress and inflammation in the central nervous system, contributing to the neuroinflammatory milieu characteristic of MS. Further investigation into the specific mechanisms and consequences of elevated HMGB1 levels could enhance our understanding of disease dynamics and possibly unveil novel therapeutic targets.

**Fig. 3.**
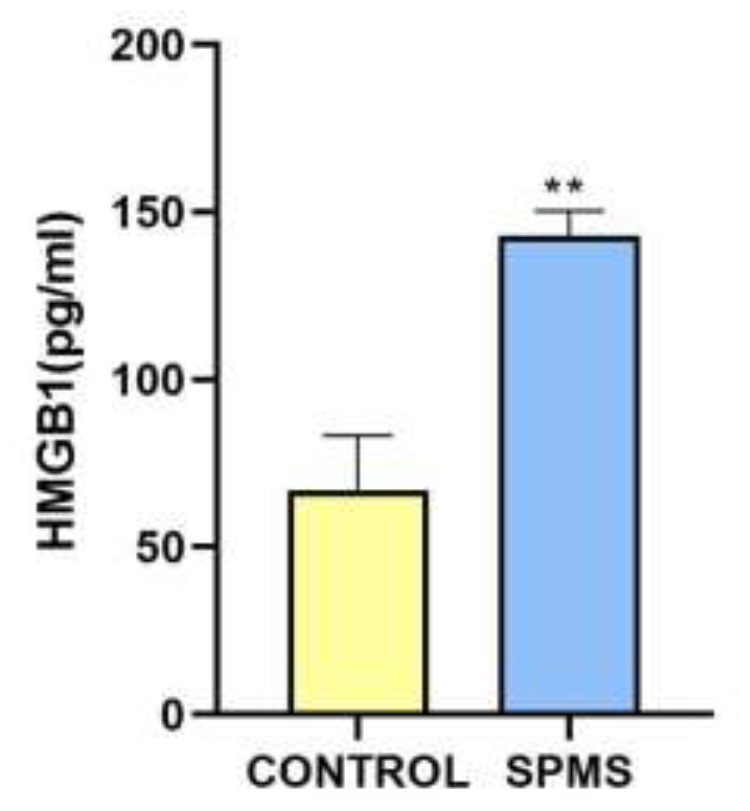
ELISA measurement of HMGB1 levels in CSF of healthy individuals and SPMS patient. ELISA assay measured HMGB1 concentrations in CSF samples of healthy individuals (control group) and SPMS patient. Each sample was analyzed in triplicate. Data were reported as mean ± SD. Significant differences between groups are indicated (**: shows values significantly different from the control group by Student’s t-test, p < 0.01).

In conclusion, this study provides valuable insights into the complex interplay between ICD and MS. Our investigation of ICD biomarkers, particularly Annexin A1, CRT, and HMGB1, in the CSF of the SPMS patient reveals significant elevations in Annexin A1, CRT, and HMGB1 levels compared to those without MS. The heightened levels of these biomarkers suggest their potential roles in the neuroinflammatory responses characteristic of MS. These findings contribute to our understanding of MS pathophysiology and may pave the way for novel diagnostic and therapeutic avenues. Although continued exploration of the specific mechanisms underlying ICD biomarkers’ variations is crucial for advancing our comprehension of the disease dynamics and identifying potential targets for more targeted therapeutic interventions in the realm of MS, meanwhile, our data indicate ICD occurrence in MS. Accordingly, prescribing cell damage and cell stress interfering compound could be beneficial for MS patients. Further studies from this group are in progress.

## Acknowledgments

The authors acknowledge personnel of Women Ward, Razi Hospital, Tabriz, Iran. The authors also acknowledge Mr Tabrizi and Clinical Laboratory of Nomouneh, Tabriz, Iran for providing MS samples data. Unfortunately, these data were not match with this study and were not included in this paper.

## Ethical statement

Ethical consent was approved by an ethics committee at Tabriz University of Medical Sciences, Tabriz, Iran. Ethic Code No: IR.TBZMED.REC.1402.348.

## Authors’ Contributions

Conceptualization, Project Administration, Supervision: Mohammad Saeid Hejazi.

Patients Selection & Samples Collection: Masoud Nikanfar, Mohammad Saeid Hejazi, Mahnaz Talebi, Vahid Hoseini.

Performing Experiments: Sevda Jafari, Mohammad Saeid Hejazi, Soheila Montazersaheb.

Data Validation, Formal Analysis: Soheila Montazersaheb, Mohammad Saeid Hejazi.

Writing, Review & Editing: Sevda Jafari, Soheila Montazersaheb, Ommoleila Molavi,

Mohammad Saeid Hejazi.

## Study Highlights

### 1) What is the current knowledge?

✓ DAMPs are considered as danger signals that elicit the innate and adaptive immune responses to enhance overt autoimmune disorder.

### 2) What is new here?

✓ The levels of Annexin A1, CRT, and HMGB1, as DAMPs, significantly elevated in CSF sample of SPMS patient compared to those without MS.
✓ The study provides valuable insights into the complex interplay between [immunogenic cell death](https://onlinelibrary.wiley.com/doi/abs/10.1111/imr.13269) and multiple sclerosis.

